# SARS-CoV-2 Subunit Virus-Like Vaccine Demonstrates High Safety Profile and Protective Efficacy: Preclinical Study

**DOI:** 10.1101/2022.05.18.492452

**Authors:** A.V. Vakhrusheva, A.V. Kudriavtsev, N.A. Kryuchkov, R.V. Deev, M.E. Frolova, K.A. Blagodatskikh, M. Djonovic, A.A. Nedorubov, E. Odintsova, A.V. Ivanov, E.A. Romanovskaya-Romanko, M.A. Stukova, A.A. Isaev, I.V. Krasilnikov

## Abstract

Public health threat coming from a rapidly developing COVID-19 pandemic calls for developing safe and effective vaccines with innovative designs. This paper presents preclinical trial results of “Betuvax-CoV-2”, a vaccine developed as a subunit vaccine containing a recombinant RBD-Fc fusion protein and betulin-based spherical virus-like nanoparticles as an adjuvant (“Betuspheres”). The aim of the study was to demonstrate vaccine safety in mice, rats, and Chinchilla rabbits through acute, subchronic, and reproductive toxicity studies. Along with safety, the vaccine demonstrated protective efficacy through SARS-CoV-2-neutralizing antibody production in mice, rats, hamsters, rabbits, and primates (rhesus macaque), and lung damage and infection protection in hamsters and rhesus macaque model. Eventually, “Betuvax-CoV-2” was proved to confer superior efficacy and protection against the SARS-CoV-2 in preclinical studies. Based on the above results, the vaccine was enabled to enter clinical trials that are currently underway.

## Introduction

Nowadays, the world population is struggling with an outbreak of the coronavirus disease-19 (COVID-19), caused by the severe acute respiratory syndrome-related coronavirus 2 (SARS-CoV-2) [1]. The World Health Organization (WHO) reported over 400 million confirmed cases of the COVID-19 and up to 6 million deaths by February 8, 2022 [2]. As a result, much effort was dedicated towards developing safe and efficient vaccines. By February 7, 2022, there were 183 vaccine candidates, and only 10 of them were approved for use by the WHO [3]. Nevertheless, by mid-February 2022 only 61.9% of the world population received at least one dose of a vaccine against COVID-19, that being approximately 10 billion administered doses, but mostly in high-income countries, whereas in low-income countries only 10.6% [4]. Moreover, fast evolution of the SARS-CoV-2 and the emergence of new contagious strains like Omicron, confirms the urgent need for the development of safe, effective, and easy scalable vaccines, including those for booster vaccination.

The Spike (S) protein of the SARS-CoV-2 is a major component that is required for the receptor binding, membrane fusion, and viral penetration. The S protein is processed at the S1/S2 cleavage site by host cell proteases, thus generating the N-terminal S1-ectodomain and a C-terminal S2-membrane-anchored protein. The S1 domain is consisted of the following subdomains: the NTD (N-terminal domain), the RBD (Receptor-Binding-Domain; residues 319-527), the S1 and S2 subdomains (SD1, SD2) [5]. The RBD is responsible for the interaction with angiotensin-converting enzyme 2 (ACE2), the primary SARS-CoV-2 receptor that is expressed in many tissues, including type II alveolar epithelial cells [6]. Therefore, the RBD is an antigen eliciting neutralizing antibodies, and thus the major target in vaccine development.

We have developed a subunit virus-like vaccine “Betuvax-CoV-2” against the SARS-CoV-2 based on the RBD and the SD1 domain of the S1 protein, fused with the human IgG1 Fc-fragment. This coronavirus antigen is being absorbed on the surface of betulin-based spherical particles 100–180 nm in size, mimicking the SARS-CoV-2 [7]. In the present paper, we have summarized the preclinical findings of the “Betuvax-CoV-2” vaccine on its safety, immunogenicity, and protectiveness.

## Methods

### Animals & Ethics Statement

BALB/c and C57BL/6 mice used in the study were obtained from the Stolbovaya branch of the FSBI SCBT FMBA (Scientific Center of Biomedical Technologies of Federal Medical and Biological Agency) of Russia. One part of the study was conducted at the Smorodintsev Research Institute of Influenza of the Ministry of Health of the Russian Federation, Saint Petersburg. The experiments on the outbred mice, rats and rabbits were carried out at the Federal State Autonomous Educational Institution of Higher Education I.M. Sechenov First Moscow State Medical University of the Ministry of Health of the Russian Federation, Moscow, Russia. Golden hamsters used in the immunogenicity and safety studies were obtained from the Branch of the Shemyakin–Ovchinnikov Institute of Bioorganic Chemistry of the Russian Academy of Sciences (Nursery for the Laboratory Animals, Puschino, Russia). The experiments on rhesus monkeys were conducted at the Research Institute of Medical Primatology, Sochi, Russia. The animals were infected at the FGBU (Federal State Budgetary Institution) Central Research Institute No. 48 of the Russian Ministry of Defense, Sergiev Posad, Russia.

The number of the animals used in the study was sufficient to assess safety, immunogenicity and protective properties of the vaccine, and at the same time, the number was sufficient to comply to the principles of ethical research. All animals were kept under standard conditions in accordance with the Directive 2010/63/EU of the European Parliament and the Council of the European Union of September 22, 2010, on the protection of animals used for scientific purposes. Animal husbandry was compliant with each facility’s SOPs and sanitary and epidemiological rules SR 2.2.1.3218-14 “Sanitary and Epidemiological Requirements for the Device, Equipment and Maintenance of Experimental Biological Clinics (Vivariums)” approved by the Resolution of the Chief State Sanitary Physician of the Russian Federation of August 29, 2014, No. 51.

Animals were kept in autoclaved sterile polycarbonate cages with Lignocel Wood Fibers wood pellets (JRC, Rosenberg, Germany) as bedding and fed *ad libitum* using standard food and purified water, under the following conditions: 20–24°C, relative humidity range of 45–65%, and light/dark cycle of approximately 12 hours per day.

The study was conducted in accordance with ARRIVE guidelines.

### Safety Studies

#### Acute toxicity study

An acute toxicity study has been performed on the outbred mice (*n*=120, 60M and 60F) and outbred rats (*n*=120, 60M and 60F). The animals were divided into groups according to their body weight (the requirement was that the difference between the individual body weight and the mean is less than 10%). Both mice and rats were administered the vaccine intravenously or intramuscularly once at a dose of 10 μg or 40 μg or placebo (*n*=20 per group, 10M and 10F). The body weight of the animals was determined at the beginning of the experiment (as previously indicated), as well as on the 7^th^ and the 14^th^ day after the administration. Clinical signs of the animals were recorded daily. Animals were sacrificed on the 14^th^ day of the experiment.

#### Subchronic toxicity study

Subchronic toxicity study of the vaccine was carried out in outbred rats (*n*=60, 30M and 30F) and Chinchilla rabbits (*n*=24, 12M and 12F). Both groups of animals were administered the vaccine intramuscularly at a dose of 5 or 20 μg or placebo (rats: *n*=20 per group, 10M and 10F; rabbits: *n*=8 per group, 4M and 4F). The administrations were performed once every 10 days over a 30-day period (4 injections in total). Condition and behavior of the animals were monitored daily, as well as the body weight and the presence of symptoms of intoxication. On the 31^st^ and the 44^th^ day after the first administration, general and biochemical blood analyses, coagulometry, urinalysis, and behavioral reactions were performed. Animals were sacrificed on the 31^st^ and the 44^th^ day of the experiment.

#### Reproductive toxicity in female rats

The reproductive toxicity of the vaccine was studied in female (*n*=60) and male (*n*=30) outbred rats. Female rats were administered the vaccine intramuscularly at a dose of 5 or 20 μg or placebo once a week over a 15-day period, before mating with males (3 injections in total, during 3 estrous cycles). A 0.9% sodium chloride solution was used as placebo. Following the administrations, females were placed together with intact males (2:1) for 10 days (2 estrous cycles) (*n*=30 per group, 10M and 20F). Condition and behavior of the animals were monitored daily. Body weight of female rats was determined before the start of the study, after the last administration of the vaccine and sacrificing on the 20^th^ day of the gestation period. The first possible day of pregnancy was determined by the presence of spermatozoa in vaginal swabs.

On the 20^th^ day of pregnancy, half of the females (*n*=30) were sacrificed for the examination of reproductive system and pre- and post-implantation mortality (the number of corpus luteum in the ovaries and the number of implantation sites, live and dead fetuses in the uterus, fetuses mass and their cranio-caudal size). The fertility index was calculated as a ratio of the number of pregnant female rats to their total number. Half of the pregnant female rats from each group were monitored until the delivery of the offspring. The offspring was then monitored for 35 days continually and evaluated on general physical condition, behavior, body weight change, and mortality.

#### Reproductive toxicity in male rats

Reproductive toxicity of the vaccine was studied in female (*n*=60) and male (*n*=30) outbred rats. Male rats were administered the vaccine intramuscularly at a dose of 5 or 20 μg or placebo once a week for 48 days before mating with females (7 injections in total). A 0.9% sodium chloride solution was used as placebo. At the end of the experiment, intact female rats were placed together with male rats for mating (1:2) for 10 days (*n*=30 per group, 10M and 20F). On the 20^th^ day of the gestation period, half of the intact female rats and all male rats were sacrificed for the examination of the reproductive system (histological and morphological analysis of the testicles, functional state of the spermatozoa and the index of spermatogenesis, in addition to the same analyses that were used in the previously described experiment). Half of the pregnant female rats were monitored until the offspring was delivered. The offspring was continually monitored for 35 days for general physical condition, behavior, body weight change, and mortality.

#### Embryo- and fetotoxicity study

Embryo- and fetotoxicity study of the vaccine was performed on mature female (*n*=60) and male rats (*n*=30). Sexually mature males were kept with females (1:2) for the duration of 2 estrous cycles (10 days) (*n*=30 per group, 10M and 20F). The first possible day of pregnancy was determined by the presence of spermatozoa in vaginal swabs. Pregnant female rats were administered the vaccine intramuscularly at a dose of 5 or 20 μg or 40 μg or placebo once a week till the 19^th^ day of the pregnancy (3 injections in total). Half of the pregnant females were monitored until the offspring was delivered and evaluated on general physical condition, behavior, and body weight changes. Three to four days before the delivery, females were each placed in a separate cage to record childbirth, the duration of the gestation period and lactation.

The number of pups, live and stillborn fetuses, and mortality was evaluated over a 35-day period post delivery. Body weight of the pups was determined at birth and on the 4^th^, 7^th^, 14^th^, 21^st^, 28^th^, and the 35^th^ day after the delivery. Cranio-caudal size was measured on the 4^th^ day. The appearance of the coat, opening of the eyes, detachment of the auricle, as well as motor activity, emotional reaction, and the ability to coordinate movements were continually monitored and recorded. The effect on the rate of maturation of sensory-motor reflexes during the feeding period were studied using the following tests: negative geotaxis, flipping on a plane, cliff avoidance, and flipping in free fall tests.

### Immunogenicity

The immunogenicity was studied in mice, golden hamsters, and Chinchilla rabbits. C57/ black mice (*n*=22) were immunized intraperitoneally twice at a dose of 5 or 20 μg or 40 μg /animal (*n*=7 per group) or PBS (*n*=8) over a 21-day period. Blood sera were taken on the 21^st^ and the 43^rd^ day of the experiment.

BALB/c mice (*n*=8) were administered the vaccine intraperitoneally twice at a dose of 5 μg or 40 μg /mouse (*n*=5) or PBS (*n*=3) over a 10-day period. Blood sera were taken on the 10^th^ and the 24^th^ day of the experiment.

Outbred mice (*n*=20 per group) and Chinchilla rabbits (*n*=8 per group) were administered two intramuscular shots at a dose of 5 or 20 μg or 40 μg or placebo (0.9% NaCl) over a 21-day period. Blood serum samples were taken from rabbits on the 21^st^ and the 35^th^ day and from the mice on the 21^st^ and the 42^nd^.

Hamsters (*n*=36) were administered the vaccine intramuscularly at a dose of 5 or 20 µg/ animal or placebo (0.9% NaCl) (*n*=12 per group, 6M and 6F), twice over a 28-day period (the 1^st^ and the 29^th^ day of the experiment). On the 36^th^ day of the experiment, blood samples from the hamsters were collected for the neutralizing antibody titers analysis.

### Anti-SARS-CoV-2 S-protein Specific IgG by ELISA

The presence of total immunoglobulins against the SARS-CoV-2 was determined using the “SARS-CoV-2-CoronaPass test system” (Biopalitra, Moscow, Russia). Antibody titers were expressed as log base 2 (Log_2_). The conversion of the logarithmic titers to the actual titers is presented in Table 1.

**Table 1.**
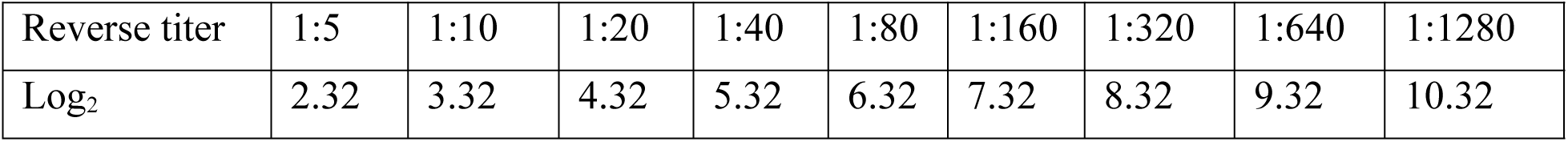
Logarithmic titer data in reverse titer

Neutralizing antibodies against the SARS-CoV-2 in hamster blood plasma were determined using the “SARS-CoV-2 Surrogate Virus Neutralization test kit” (GenScript, USA).

Enzyme immunoassays were carried out using the “Multiskan GO” microplate spectrophotometer (Thermo Fisher Scientific, Waltham, Massachusetts, United States) at 450 nm.

### Protective Efficacy

The protective efficacy of the vaccine was evaluated in a study using hamsters (*n*=72) and rhesus monkeys (*Macaca mulatta*) (*n*=18). Hamsters were administered the vaccine intramuscularly at a dose of 5 or 20 μg or 40 μg /animal (0.5 ml/animal) or intranasally at a dose of 4 μg or 40 μg (100 µl/animal, 10 µl in each nostril 5 times a day with an interval of 30 minutes). The control group received placebo (0.9% NaCl) intramuscularly or intranasally or were not administered any solution (virus dose control group) (*n*=12 per group, 6M and 6F). Healthy male rhesus monkeys (*n*=18) received 0.5 ml of the vaccine intramuscularly at a dose of 5 or 20 μg or 40 μg /animal or placebo (0.9% NaCl) (*n*=6 per group). The vaccine was administered twice on the 1^st^ and the 29^th^ day of the experiment (over a 28-day period). Following the immunization, the animals were infected intranasally with the SARS-CoV-2 at a dose of 10^5^ PFU (PFU — plaque-forming unit).

#### Infection of animals with the SARS-CoV-2

For the preparation of the infectious mixture, a SARS-CoV-2 virus isolate B strain culture was used in the form of a freeze-dried 30% Vero C1008 cells suspension in a 10% sucrose water solution with the addition of a 10% fresh chicken egg yolk. The initial biological activity of the culture was 7±0.2 lg PFU/ml. Content of two ampules containing the cell culture was resuspended in Hank’s solution supplemented with 2% FCS (fetal calf serum) (Thermo Fisher Scientific, Waltham, Massachusetts, United States). The solution was administered intranasally at a dose of 10^5^ PFU/animal.

For further experiments, lung samples were taken from 4 hamsters and 2 monkeys from each group on days 2, 4 and 6 after the infection. The SARS-CoV-2 accumulation titer in the lungs was determined using the negative colonies method on the Vero C1008 cell culture. Lung samples from macaques were used for histology and immunohistology analysis.

#### Determination of the VSI (virus suppression index) of the SARS-CoV-2 in the lungs of golden hamsters

Hamster lungs were homogenized and diluted in Hank’s solution with the addition of 2% FCS and antibiotics (streptomycin sulfate and benzylpenicillin sodium salt, 200 U/ml) (Thermo Fisher Scientific, Waltham, Massachusetts, United States) to prepare a 10% lung suspension. Biological activity of the virus from the lung samples was determined by titrating the prepared suspension to a monolayer of Vero C1008 cell culture in plastic vials with a 25 cm^2^ working surface area (CellStar, Greiner Bio-One GmbH, Kremsmünster, Austria). After removing the medium, a monolayer of Vero C1008 cells was infected with 10-fold dilutions of the virus. Vials with an infected monolayer were incubated for 1 hour at 36.5– 37.5°C. The inoculum was decanted and the primary agar coating, prepared according to the prescription for the SARS-CoV-2, was added. The cell monolayer was incubated for 2 days at 36.5-37.5°C, after which a secondary agar coating was added and incubated for 24 hours at the same temperature. After the completion of this step, the number of negative colonies was determined. A decrease in the level of viral load at the peak of infection was taken as a parameter for evaluating the vaccine efficacy (at a 95% confidence level — decrease level from 1.2 to 1.8 lg, at 99% confidence level — more than 1.8 lg).

#### Histological study

In this study, 18 lung complexes from monkeys were examined, including trachea, main bronchi, left and right lungs, esophagus, fragments of the aorta, fat tissue of the mediastinum, thymus, and lymph nodes (the sections were fixed in a 10% formalin). The material was excised with macroscopic photodocumentation. As a result, over 120 diagnostically significant areas of the lungs were obtained for histopathological micropreparations.

Using a computer video system (IBM PC + microscope Leica DM 1000, Leica Camera, Wetzlar, Germany) and the ImageScope-M software package (Systems for Microscopy and Analysis, Moscow, Russia), the “airiness” parameter, specific area of the histological section occupied by air and free from infiltration or exudate from the alveolar space, was determined in a semi-automatic mode.

An immunohistochemical study was performed to immunotype infiltrated cells in the lungs using antibodies against the T-lymphocyte marker (CD3 — Cell Marque, 103-R94, Rocklin, California, USA) and macrophages M2-phenotype (CD163 MsmAb — Cell Marque, 163M-15, Rocklin, California, USA). Antibodies against vascular endothelial cells (CD31 Mon. mouse – Cell Marque, JC-70, Rocklin, California, USA) were also used. Morphometric evaluation consisted in counting the number of cells per 1 mm^2^ area of the lung parenchyma.

For immunohistochemical evaluation of the coronavirus proteins (S- and N-proteins), SARS-CoV2 spike (Genetex, GTX135356, Irvine, California, USA) and SARS-CoV2 nucleocapsid (Invitrogen, MA1-7404, Waltham, Massachusetts, USA) antibodies were used. The result was evaluated visually and qualitatively. The lung tissue from a patient who died from a coronavirus infection served as a positive control. The negative control was the lung tissue from a patient who died from causes not related to the respiratory system.

### Statistical Analysis

The obtained data were analyzed with Microsoft Office Excel 2010 (Microsoft, Redmond, Washington, USA) and GraphPad Prism v6.01 software (GraphPad Software, San Diego, California, USA) for geometric mean, standard deviation, arithmetic mean, and standard errors. No samples or animals were excluded from the analysis. The Kruskal–Wallis H test, or its parametric equivalent one-way analysis of variance (ANOVA), was used for the analysis of variance between the group means, with subsequent pairwise comparison using Tukey test or non-parametric pairwise multiple comparisons by Dunnett’s test. Values p <0.05 were accepted as statistically significant.

## Results

### Betuvax-CoV-2 vaccine safety in toxicity studies

#### Acute and subchronic toxicity studies

The acute toxicity study in outbred mice and rats with single intravenous and intramuscular vaccine injections demonstrated that a single administration of the vaccine at 10 μg/animal or 40 μg/animal did not lead to mortality, body, or organ weight decrease, or worsening of the overall condition. The determination of lethal doses LD10, LD16, LD50, or LD84 was impossible due to absence of mortality. It was also shown that the animals steadily gained weight with no lags for the entire period of the study. The average values of the initial body weight and the body weight at the end of the study are depicted in Figure 1. Comparison of six groups (males and females separately) using the Kruskal-Wallis test did not show any significant differences (p >0.05). The necropsy study, carried out on the 14^th^ day following the vaccine administration, showed no abnormalities in the structure of the internal organs in mice and rats or any pathological changes (data not shown).

**Figure 1.**
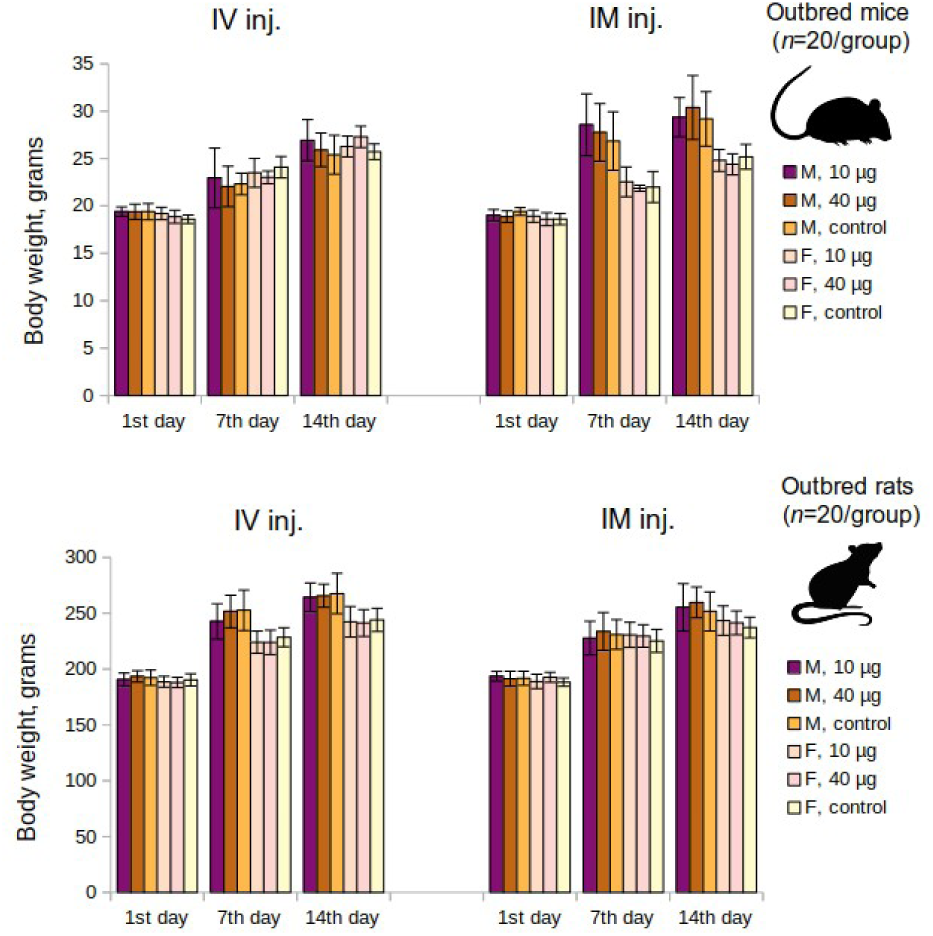
Mean changes of body weight over time in the acute toxicity study in mice and rats groups. M - males, F - females, IM inj. - intramuscular injection, IV inj. - intravenous injection; p >0.05

Subchronic toxicity studies in rats and Chinchilla rabbits showed that there were no differences observed between the experimental group and the control in terms of the activity, movement, appearance, physiological functions, animal behavior, bone marrow cellularity or myelogram parameters, body weight, based on hematological, biochemical, or coagulometric parameters and urine analysis.

The necropsy study did not reveal any deviations in the structure of the body and the internal organs in experimental groups. There were also no intergroup differences in the mass of internal organs (heart, lungs, liver, kidneys, brain) including immunocompetent organs (thymus, spleen). Histological examination of the internal organs and tissues in experimental animals and the control, demonstrated no pathological changes (data not shown).

The absence of toxicity, including the lack of immunotoxicity and local irritation, was demonstrated for the “Betuvax-CoV-2” vaccine in the acute and subchronic toxicity studies in mice, rats, and rabbits.

#### Reproductive toxicity study in female rats

To evaluate the condition of the reproductive system of pregnant female rats, immunization was performed 15 days before mating. The administration of the “Betuvax-CoV-2” vaccine at doses of 5 and 20 µg/animal did not affect the reproductive function adversely (Figure 2). The embryo mortality at all stages of the development in female rats immunized with the vaccine did not differ from the control. There were also no significant differences in fetus weight or size (p >0.05, Kruskal-Wallis test).

**Figure 2.**
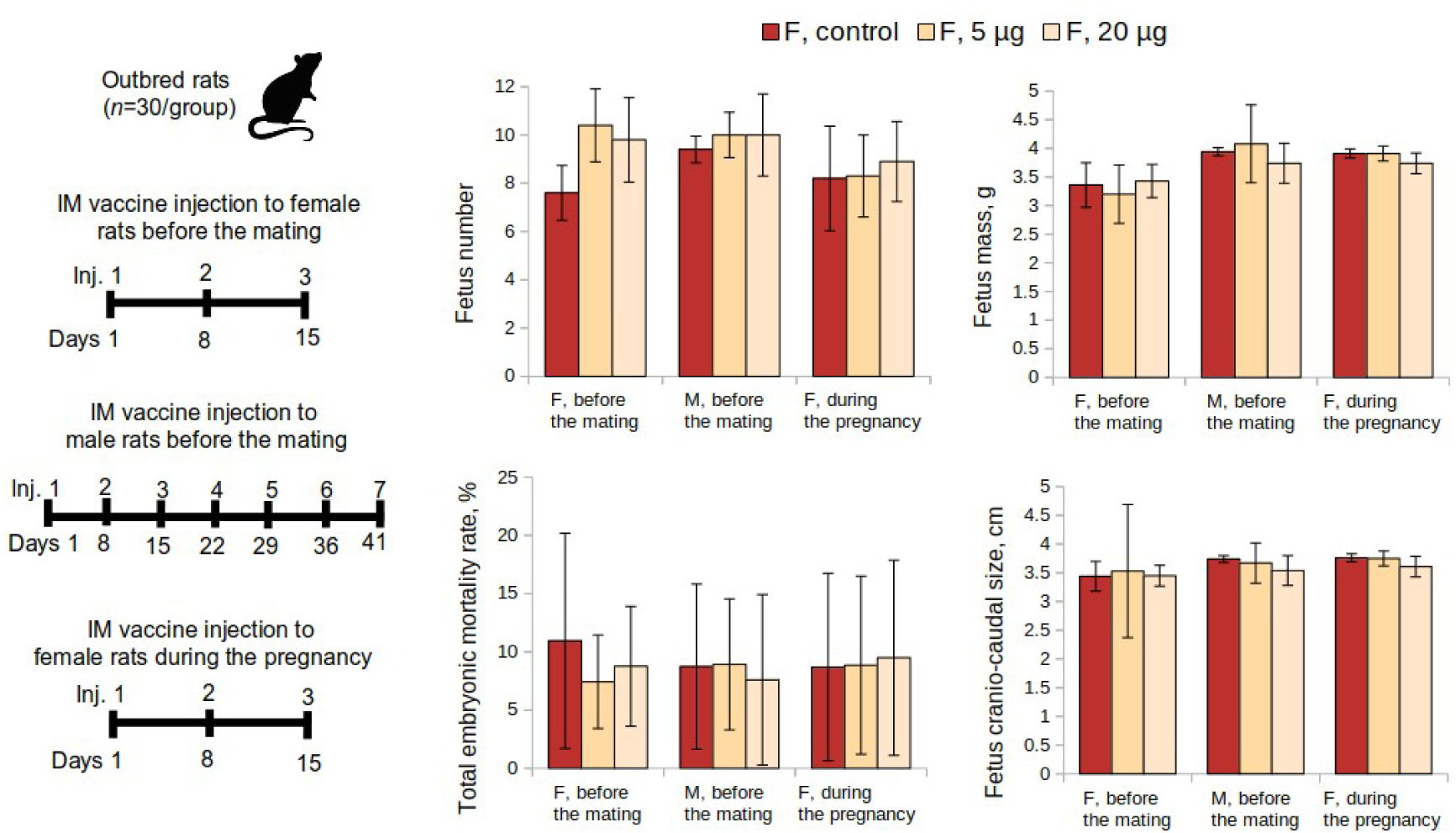
Examination of the reproductive system according to the following parameters: fetus number, fetus mass, total embryo mortality rate, fetus cranio-caudal size from female rats intramuscularly immunized with the “Betuvax-CoV-2” vaccine 15 days before mating (F, before the mating), before mating with males previously immunized during a 48-day period (M, before the mating), and during the first 19 days of the pregnancy (F, during the pregnancy). F - females, M - males; IM - intramuscular administration; p >0.05

Macroscopic and histological examination of rats’ ovaries that were immunized with the “Betuvax-CoV-2” vaccine within 15 days before mating showed no signs of pathological changes. The ovaries in all groups were dark red, moderately dense and with an uneven surface. Follicles of various sizes and degrees of maturation were visible. The follicular epithelium was not changed; the nuclei were light, clear, and the ovarian medulla was full-blooded (data not shown).

The effect of the “Betuvax-CoV-2” vaccine when administered to female rats 15 days before mating was assessed on the postnatal development of the offspring (Figure 3). Half of the pregnant rats were monitored continually until the delivery of the offspring. The development of the offspring was observed for 35 days for general physical condition, behavior, weight dynamics, and mortality. All pregnant animals of the experimental groups demonstrated normal birth and labour processes. The general and specific indicators of postnatal development did not differ significantly from the control values (p >0.05, Kruskal-Wallis test). The duration of the pregnancy in experimental females was in accordance with the physiological norm for this type of animal and the control. All rat pups were born viable and no mortality during the first month was recorded. The physical development of the offspring proceeded without any deviations.

**Figure 3.**
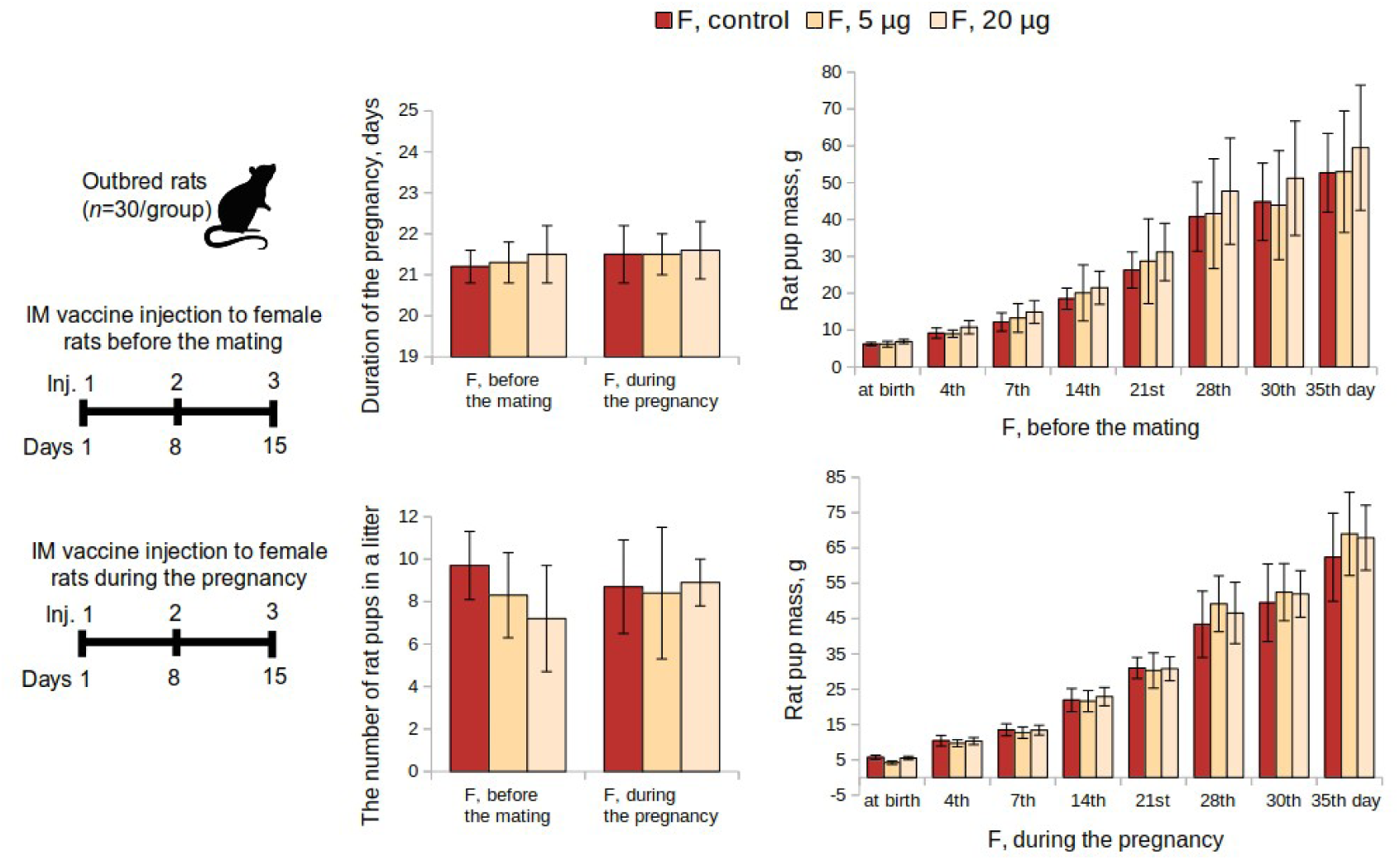
Evaluation of postnatal development of the offspring according to the following parameters: duration of the pregnancy, the number of rat pups in a litter, rat pup mass, when female rats were intramuscularly immunized with the “Betuvax-CoV-2” vaccine 15 days before mating (F, before the mating); during the first 19 days of the pregnancy (F, during the pregnancy). F - females; IM - intramuscular injection; p >0.05

#### Reproductive toxicity study in male rats

The condition of the reproductive system of the pregnant rats who were mated with the males who were previously administered the “Betuvax-CoV-2” vaccine for 48 days before mating (at doses 5 μg and 20 μg) is depicted on Figure 2. The values indicating the condition of the reproductive system did not differ from those females mated with the males of the control group (p >0.05, Kruskal-Wallis test). In this way, the administration of the vaccine did not adversely affect the reproductive function of male rats.

Macroscopic testicle study in outbred male rats that were intramuscularly administered the “Betuvax-CoV-2” vaccine at doses 5 μg and 20 μg once a week for 48 days demonstrated no pathological changes. Testicles of all experimental groups were of unchanged compared to the control, oval, pale gray in color; the blood vessels with well expressed lobulated structure (data not shown).

Histological study in rat testicles showed that the administration of the vaccine at 20 μg or 40 μg or placebo μg once a week for 48 days had an adverse effect on the condition of the spermatogenic epithelium expressed in abnormal polarity and germ cells localization. It manifested as cell desquamation into the lumen of seminiferous tubules (in 6 out of 10 males of the 20 μg or 40 μg group). Spermatogenesis Index of the rats administered the vaccine at 5 μg or 40 μg did not show significant intergroup differences (p >0.05, Dunnett’s test). However, the 20 µg group was significantly different from the control (p <0.001, Dunnett’s test), the absolute difference between the sample means in the control group and the 20 µg group was only 0.22. Thus, the spermatogenesis in the 20 μg or 40 μg group was decreased slightly and had not pronounced effect on reproductive potential of male rats.

Statistically significant intergroup differences in the functional state of the spermatozoa of rats that were administered the “Betuvax-CoV-2” vaccine at 5 and 20 μg or 40 μg once a week for 48 days were also not detected. There were no adverse effects on the embryo development recorded. This was confirmed by examining the reproductive system of pregnant females and the development of the offspring (Figure 2).

#### Embryo- and fetotoxicity study

Embryo- and fetotoxic effects of the vaccine were examined during the antenatal period in cases when the vaccine was administered intramuscularly to female rats during the first 19 days of the pregnancy. Intramuscular injections were administered once a week from the 1^st^ to the 19^th^ day of pregnancy at 5 or 20 μg or 40 μg and did not show embryotoxic and teratogenic effects in terms of the offspring development. Mortality was absent and there were no signs of toxicosis, visible pregnancy abnormalities, or decreased body weight gain of female rats in the experimental group in comparison to the control. Embryo mortality at all stages of the development at all doses did not differ from the corresponding values of the control group (0%). Significant differences from the control in weight and size of the fetus and placenta were also not revealed (Figure 2). No external abnormalities of the embryos were found. The study of nine sagittal sections confirmed the absence of internal developmental anomalies. No anomalies in the skeleton development and delays in the ossification of the skeleton of the embryos were also found (data not shown).

The effects of intramuscular administration of the vaccine to female rats during the first 19 days of pregnancy were evaluated during postnatal development of the offspring as well (Figure 3). Integral and specific indicators of postnatal development of rat pups from experimental female rats did not differ significantly from the control values (p >0.05, Kruskal-Wallis test). None of the doses affected the physical development of the offspring, the development of sensory-motor reflexes during the period of feeding, motor activity or emotional reactions after the end of feeding.

Physical development of the offspring was normal. The recorded time of the manifestation of specific developmental events, such as the detachment of the auricles, the appearance of the primary hairline, or the opening of the eyes were within the control and standard fluctuations for this animal species. Embryos with developmental anomalies, both external and internal, were absent from all experimental groups.

Thus, the results of the preclinical study indicate a high safety, absence of general toxicity, or reproductive toxicity of the “Betuvax-CoV-2” vaccine in mice, rats, and Chinchilla rabbits of both sexes.

### Betuvax-CoV-2 vaccine stimulates humoral immune response

#### Antibody titers in the sera of C57/black and BALB/c mice

Immunization and sample acquisition for this study were performed as shown in Figure 4, top panel. The study showed that the vaccine antigen specific antibody level has increased after a single immunization with “Betuvax-CoV-2”, yet it was not considerable. A significant increase in specific antibody titers in the immunized C57/black mice in comparison with control was observed after two intraperitoneal injections of the vaccine preparation. The IgG titers significantly increased in both groups: upon administration of the vaccine at a dose of 5 µg/animal (GMT (geometric mean titer) =1600) and at a dose of 20 µg/animal (GMT=2377), compared to control (p <0.05, Tukey’s test) (Figure 4).

**Figure 4.**
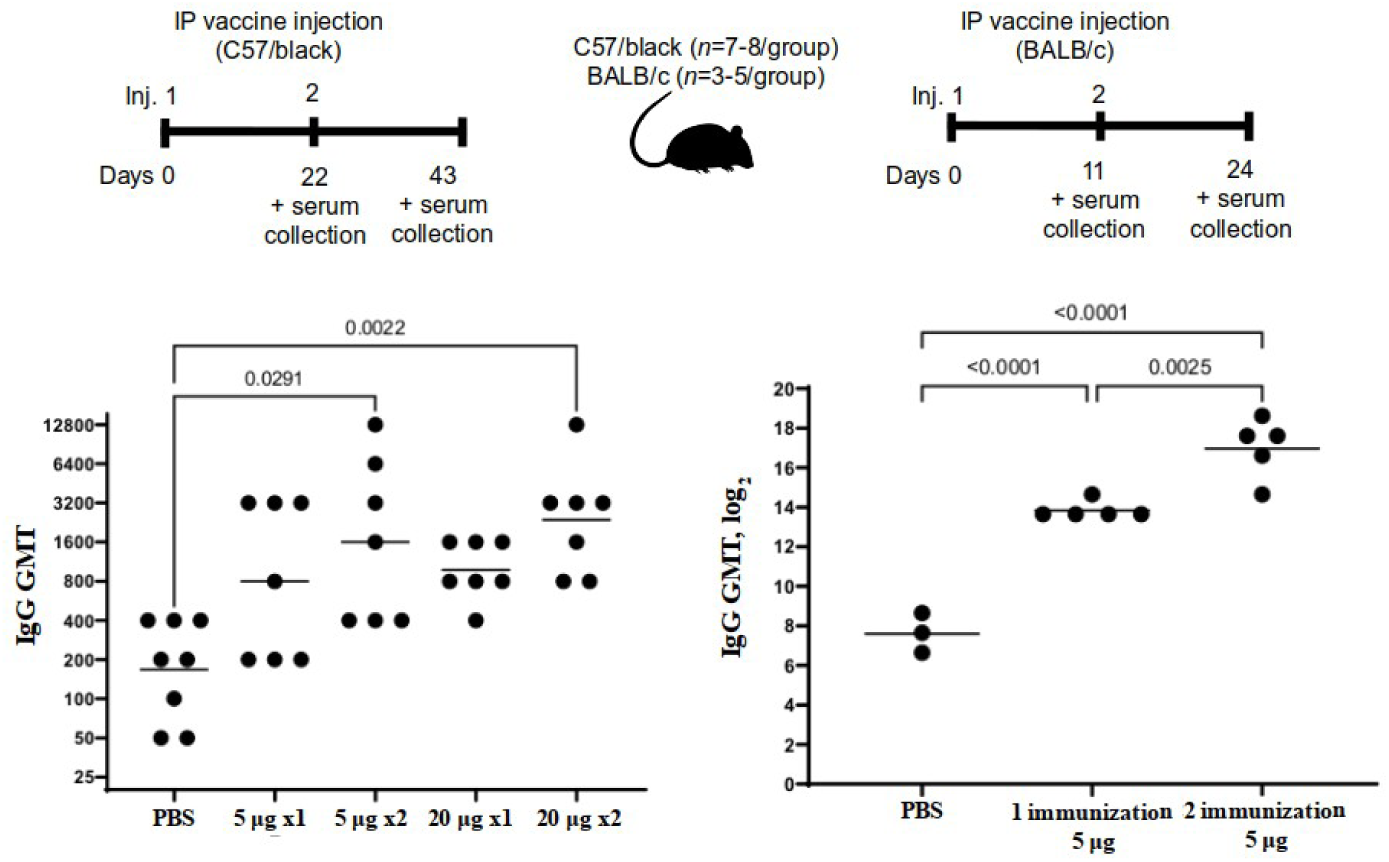
The level of antibodies against the SARS-CoV-2 RBD-based protein in the sera of immunized C57/black (left) and BALB/c (right) mice after a single (x1) or double (x2) intraperitoneal immunization with “Betuvax-CoV-2” at a dose of 5 μg or 20 μg (left) and 5 μgg or 20 μg or 20 μg (left) and 5 μgg (left) and 5 μg or 20 μg (left) and 5 μgg (right). Separate symbols (dots) represent individual values of titers in animals, the horizontal line shows average group geometric mean titers; p <0.05

A significant increase in specific antibody titers in the serum of BALB/c mice was registered after a single immunization compared to the control group (PBS). The GMT value after a single immunization at a dose of just 5 μg or 40 μg was 14703. The antibody titers after re-vaccination compared with those after a single immunization was multiplied by 9 times (GMT=132578). All groups were significantly different from each other (p <0.05, Tukey’s test) (Figure 4).

#### Antibody titers in the sera of rabbits and outbred mice

The study in rabbits and mice were performed according to the scheme on Figure 5. The study on rabbits showed that the RBD-specific antibody level after the first immunization in the 5 μg or 40 μg, 20 μg or 40 μg and the control group did not differ significantly (p=0.999, Kruskal-Wallis test), yet in the outbred mice it significantly differed (p <0.001, Kruskal-Wallis test) in the 20 µg group (Log_2_=4.40, GMT=1:21) (p <0.001, Dunnett’s test).

**Figure 5.**
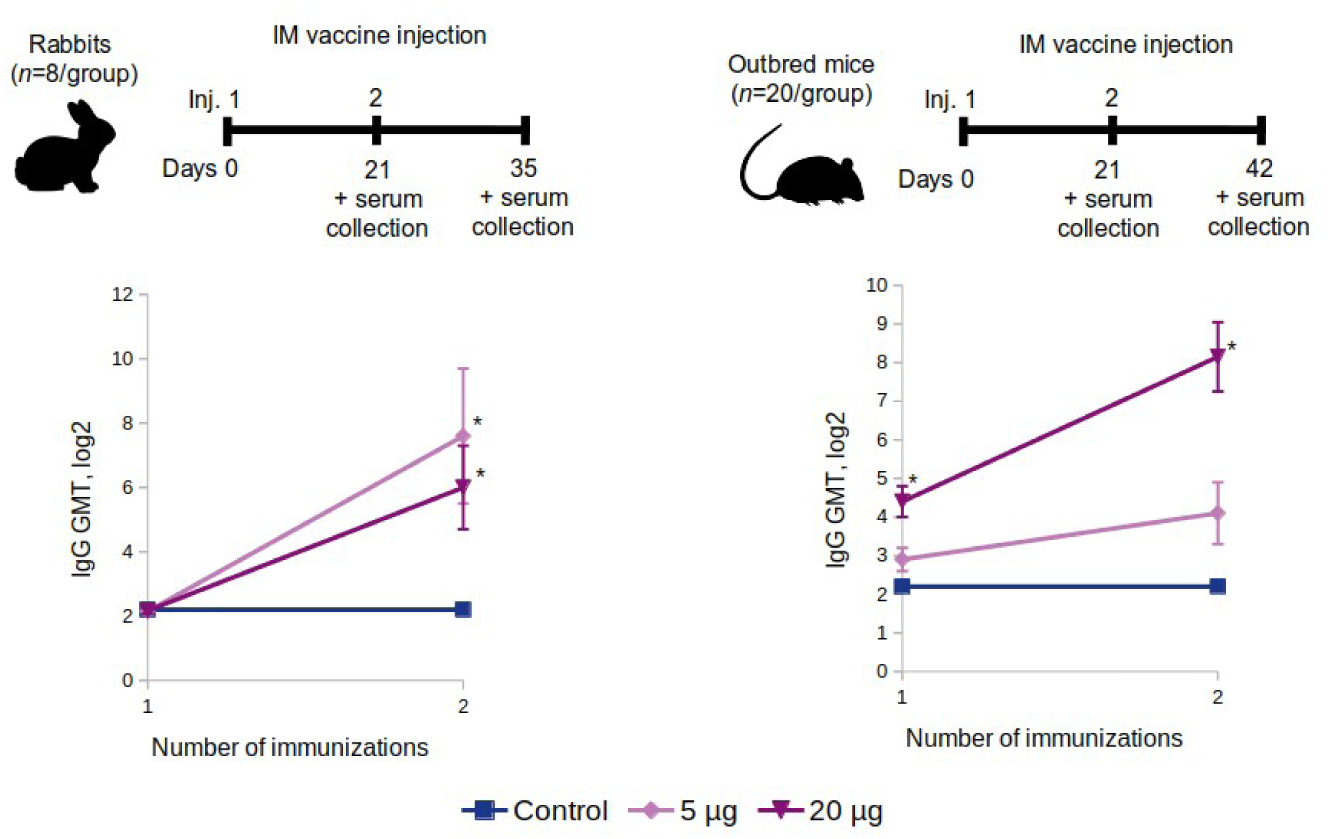
The growth of the GMT level of antibodies specific to the SARS-CoV-2 RBD protein in the serum of the intramuscularly immunized rabbits (left) and outbred mice (right) with “Betuvax-CoV-2” at 5 and 20 µg/animal; * p <0.05

In rabbits, 14 days after the second immunization (on the 35^th^ day of the experiment), the antibody level in the 5 μg or 40 μg, 20 μg or 40 μg and the control group differed significantly (p=0.044, Kruskal-Wallis test). The level of antibodies increased in the 5 μg or 40 μg group (Log_2_=7.57, GMT=1:190) (p=0.024, Dunnett’s test) as well as in the 20 μg or 40 μg group (Log_2_=6.07, GMT=1:67) (p=0.04, Dunnett’s test) compared with the control (Figure 5).

In the outbred mice, 21 days after the second immunization, on the 42^nd^ day after the first administration of the vaccine, the GMT of antibodies in the 5 µg, 20 µg and control group was significantly different (p <0.001, Kruskal-Wallis test). In the 5 μg or 40 μg group, the level of antibodies did not significantly increase (Log_2_=3.99, GMT=1:16) (p=0.088, Dunnett’s test) compared to the control group. In the 20 μg or 40 μg group, the antibody level significantly increased (Log_2_=8.24, GMT=1:305) (p <0.001, Dunnett’s test) compared to the control (Figure 5).

#### The level of neutralizing antibodies in the sera of golden hamsters

The presence of neutralizing antibodies against the SARS-CoV-2 was determined by “SARS-CoV-2 Surrogate Virus Neutralization Test Kit” (GenScript, USA) in the blood plasma of hamsters taken on 36^th^ day after two immunizations. The first group of hamsters with an intramuscular placebo injection (0.9% NaCl) showed the absence of the antibodies against the SARS-CoV-2. The second group of hamsters, immunized intramuscularly with “Betuvax-CoV-2” at 5 µg/animal, showed a response in the production of neutralizing antibodies in 33.3% of the animals (2 out of 6 females and 2 out of 6 males). The third group of hamsters immunized intramuscularly at 20 μg or 40 μg /animal produced neutralizing antibodies in 92% of cases (5 out of 6 males and 6 out of 6 females) (Table 2).

**Table 2.** Golden hamsters with neutralizing antibodies against SARS-CoV-2

| | Control<br>0.9% NaCl<br>$n=12$ (6M, 6F) | Betuvax-CoV-2<br>5 µg/animal<br>$n=12$ (6M, 6F) | Betuvax-CoV-2<br>20 µg/animal<br>$n=12$ (6M, 6F) |
| --- | --- | --- | --- |
| Hamsters with neutralizing antibodies/all hamsters | 0/12 | 4/12 | 11/12 |

To conclude on the immunogenic studies shown above, “Betuvax-CoV-2” demonstrated that all animal species (mice, rats, rabbits, hamsters) produced virus-neutralizing antibodies against the RBD component of the SARS-CoV-2.

### Betuvax-CoV-2 vaccine shows high protective efficacy against the SARS-CoV-2

#### Accumulation of the SARS-CoV-2 virus in the lungs of golden hamsters

The specific antiviral activity of the vaccine (protective efficacy) was evaluated in golden hamsters at 5 and 20 μg or 40 μg (intramuscularly) and 4 μg or 40 μg (intranasally) before the infection with SARS-CoV-2. The experiment was performed on hamsters as described on Figure 6, top panel. Neither in males nor females there were clinical signs of deviations of health condition, local irritation, or statistically significant intergroup differences in terms of body weight or body temperature during immunization period (data not shown).

**Figure 6.**
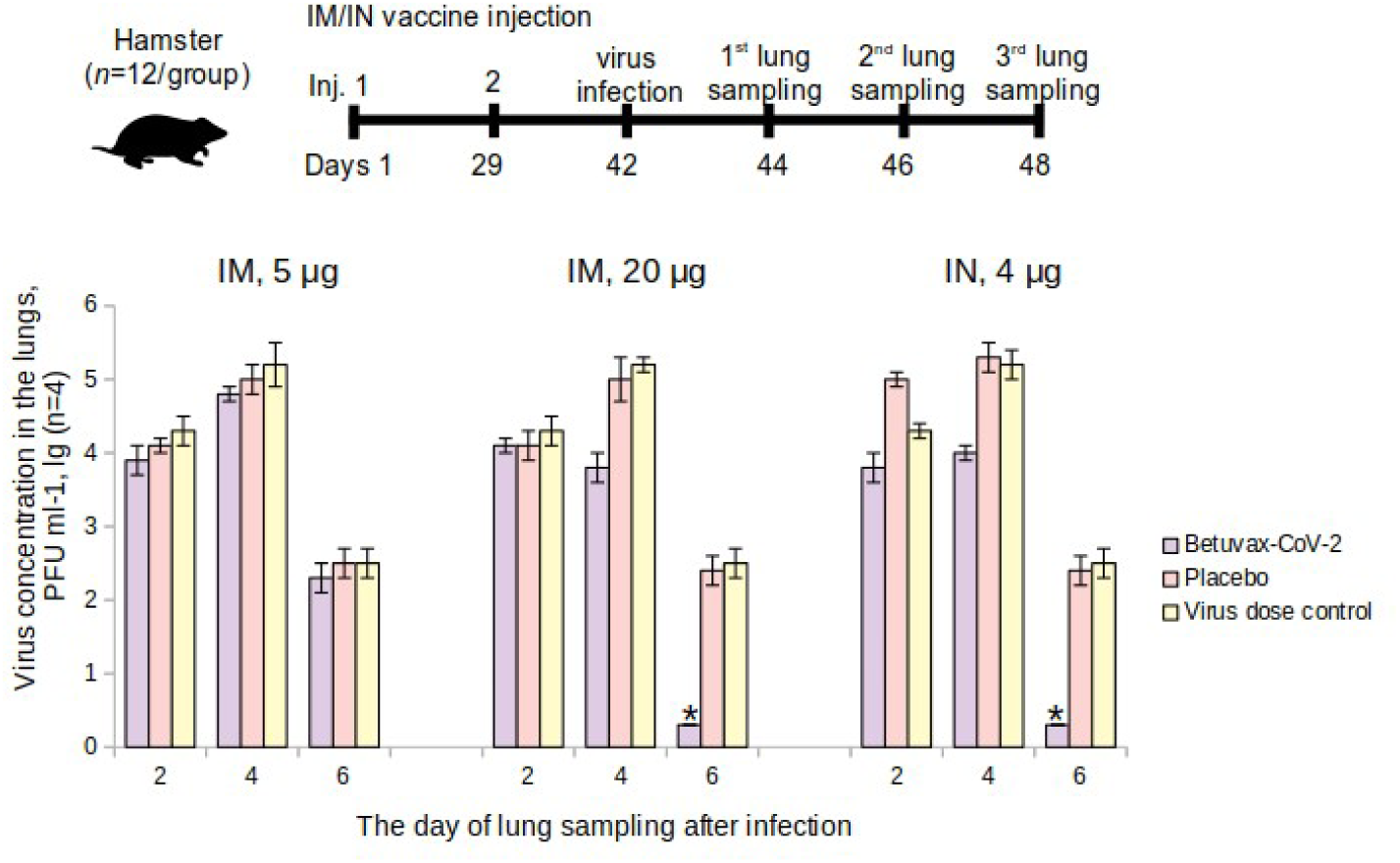
The level of the SARS-CoV-2 accumulation in the lungs of vaccinated golden hamsters on the 2^nd^, 4^th^, and the 6^th^ day after infection. IM - intramuscular, IN – intranasal administration; * p <0.05

After an intranasal infection with the SARS-CoV-2 at 10^5^ PFU in the 20 μg or 40 μg /animal hamster group (double intramuscular injection) and in the 4 μg or 40 μg /animal group (double intranasal injection) the virus concentration in the lungs was completely suppressed on the 6^th^ day of the study in comparison with placebo (Figure 6).

The virus suppression index (VSI) in the lungs of the 20 μg or 40 μg /animal hamster group was: 1.4±0.2 lg PFU·ml^-1^ — on the 4^th^ day after the infection, 2.2±0.2 lg PFU·ml^-1^ — on the 6^th^ day after infection. In the group that was immunized intranasally at a dose of 4 µg/animal (100 µl of a dosage of 40 µg/ml): 1.3±0.2 lg PFU·ml^-1^ and 2.2±0.2 lg PFU·ml^-1^ on the 4^th^ and 6^th^ day, respectively. When the hamsters were immunized intramuscularly at a dose of 5 µg/animal, the VSI ranged from 0.2±0.3 to 0.4±0.4 lg PFU·ml^-1^. In accordance with the Guidelines for Preclinical Studies, the drug is effective if the VSI is from 1.2 to 1.8 lg (95% confidence level), at 1.8 lg or more (99% confidence level). The VSI was above 1.8 lg on the 6^th^ day in the intramuscular 20 μg or 40 μg administration and intranasal 4 μg or 40 μg administration groups that testifies a complete absence of the virus production in the lungs.

Thus, it has been shown that the protective activity of a double intramuscular administration of the vaccine at 20 μg or 40 μg and the intranasal administration at 4 μg or 40 μg against the SARS-CoV-2 was 99%.

Since the level of the SARS-CoV-2 accumulation in the lungs of the immunized hamsters was lower than in the control group, it might also suggest that there was no ADE (antibody-dependent enhancement) effect.

#### Evaluation of the protective activity of the vaccine against the SARS-CoV-2 in the lungs of rhesus monkeys

The protective activity of the vaccine was studied in intramuscularly immunized rhesus macaques at 5 or 20 μg or 40 μg before the infection with the SARS-CoV-2 (Figure 7, top panel). There were no clinical signs of changes in health condition, local irritation, or significant intergroup differences in body weight or body temperature during immunization period (data not shown).

**Figure 7.**
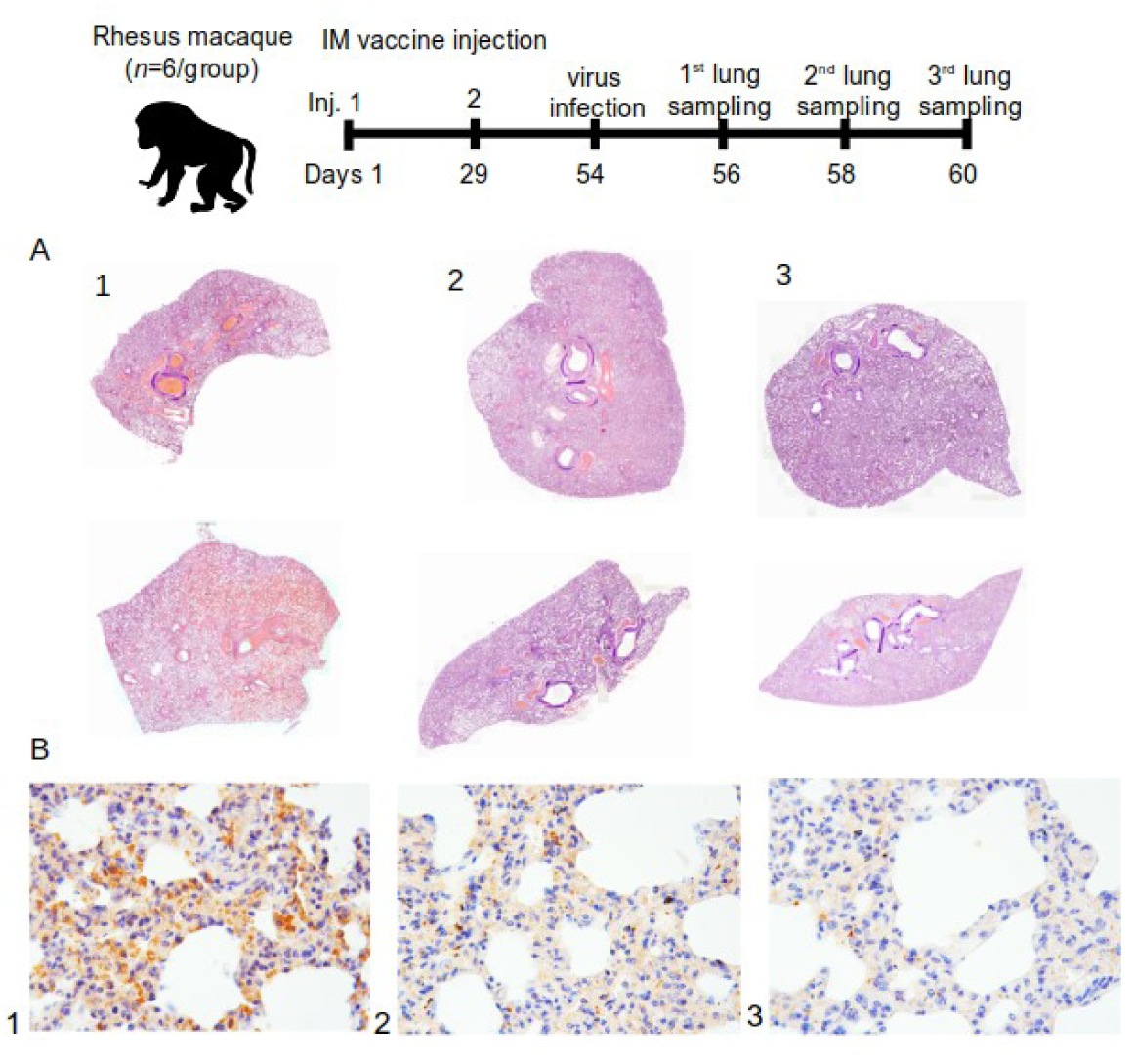
A) Histotopogram of macaque representative lung samples taken on the 6^th^ day after the infection of rhesus macaques with the SARS-CoV-2; B) IHC-reaction to S-protein (brown reaction product) in representative lung tissues of rhesus macaque (1 — control, 2 — 5 μg or 20 μg (left) and 5 μgg dose, 3 — 20 μg or 20 μg (left) and 5 μgg dose); stained with hematoxylin; magnification×250 (n=72)

After the macaques were intranasally infected with the SARS-CoV-2 at 10^5^ PFU, there were changes in pathohistology such as interstitial inflammation, widespread (in a pulmonary circulation) thrombosis (thrombovasculitis) of large, medium, and small vessels in the lung tissues. However, the tissue changes in the animals of the control group (1) were significantly different from those in the vaccine groups of 5 µg and 20 µg (2 and 3). In the control group, the pathomorphological picture showed signs of exudative inflammation and catarrhal changes in the bronchi. The most characteristic symptom was thrombo-hemorrhagic changes. In the vaccine groups, there was a slight hemorrhagic component, however, with a more pronounced manifestation of interstitial inflammation in the form of lympho-macrophage infiltration. Also, in these groups, there were no progressive lesions, necrosis, abscess formation, or signs of fibrinous effusion in the alveoli (Figure 7A).

The airiness index expresses the specific area in the lumen of alveoli on a histological section participating in external respiration and gas exchange. According to the airiness index, the number of differences found between the groups, such as signs of damage, were probably of viral origin. In fact, the presence of normal airiness foci was registered only in the lungs of animals of the vaccine groups (Figure 7A).

Tracheal injury included less pronounced signs of partial desquamation of the epithelium at the level of the lower third, as well as a minimally pronounced parietal accumulation of mucofibrinous exudate (in the control group). The animals in the vaccine groups observed no changes in the trachea for the duration of the study.

#### Immunohistochemical and morphometric characteristics of the inflammatory infiltrate

Immunohistochemical evaluation of the tissue samples was carried out using anti-human antibodies against CD163, a marker of macrophages (M2, histiocytes) and antibodies against CD3, a panlymphocyte marker. Lung tissue from a deceased person confirmed to be infected with the novel coronavirus infection was used as a positive control.

The lymphocyte-macrophage ratio was the highest in the control group on the 6^th^ day - of about 70% (66.7%). Therefore, about two thirds of the leukocytes in the infiltrate were T-lymphocytes. The most pronounced immune response at the level of tissue and according to the infiltrate characteristics was in the control group. More pronounced inflammatory manifestations were specific for the animals in the experimental groups, and the maximum number of macrophages was in the 20 μg or 40 μg group.

As it is shown in Figure 7A, lung samples from the control group (1, Figure 7A) were consistent with a subtotal hemorrhagic interstitial pneumonia. Segmental bronchi in the distal sections were filled with blood, full with fibrin and many leukocytes. All vessels were sharply plethoric, mostly thrombosed. Subpleural vessels were especially pronounced with leukocyte infiltration. There were also some foci of effusion pleurisy. In Figure 7A on the upper part of the control group (1, Figure 7A), can be seen that the lobar bronchus was entirely filled with blood. Distelectasis, thickening of the interalveolar septa due to edema and massive leukocyte infiltration, and subtotal thrombosis of the vessels of all calibers were recorded as well.

For the group with a dose of 5 μg or 40 μg (2, Figure 7A) in both samples the lobar and segmental bronchi were passable, there was no secret or exudate. Focal interstitial edema, infiltration, and partial thrombosis of large and a part of small vessels was also observed. In both samples (2, Figure 7A), areas of high airiness were also determined.

The group immunized with 20 μg or 40 μg (3, Figure 7A) was characterized with passable lobar and segmental bronchi and the absence of exudate. Peribronchial focal lympho-macrophage infiltration and focal changes were absent. Some blood vessels were either thrombosed or contained sludge, but no hemorrhagic component was observed. On the upper part (3, Figure 7A), the stroma of the organ was diffusely infiltrated with leukocytes; there were no segmented leukocytes or focal changes. In these groups (3, Figure 7A), large areas of high airiness and single lymphoid nodules were also observed.

#### Detection of viral proteins in lung tissue

The mosaic arrangement of lung tissue areas with virus S-protein was determined by immunohistochemistry (IHC) on the 2^nd^, 4^th^, and 6^th^ day after infection. Reactions with antibodies against the N-protein turned out to be negative both with the experimental material and with the control samples. Figure 7B shows that there was a trend towards a decrease in the detection of the S-protein in the lungs of primates depending on the use of the “Betuvax-CoV-2” vaccine and its dosage. The detectable protein inclusions of the S-protein in groups 5 and 20 µg (2 and 3, respectively, Figure 7B) were significantly less compared to the control group (1, Figure 7B).

Also, the ADE effect was evaluated according to the criteria of overall morphological and physiological state, as well as macroscopic changes in the lungs of the vaccinated and the control animals. Macroscopic assessment of the lungs of monkeys, as well as the immunohistochemical analysis of the accumulation of the S-protein of the SARS-CoV-2 in the lungs of the vaccinated animals, showed no ADE effect, supporting our previous findings in hamsters.

As a result of determining the protective activity of the “Betuvax-CoV-2” vaccine in intramuscularly immunized hamsters at a dose of 20 μg or 40 μg /animal and intranasally at a dose of 4 μg or 40 μg / animal, an elimination of the virus from the lungs was shown to be 99%. Histology of rhesus monkey tissue also showed that animals vaccinated intramuscularly at 5 or 20 μg or 40 μg did not demonstrate pronounced signs of damage of the vascular component, unlike the control group, but reflected an inflammatory component of the immune system to a greater extent. Also, IHC identification of the S-protein showed a significantly lower presence in the vaccinated macaques. In the study of the protection activity in both hamsters and rhesus monkeys, no manifestations of the ADE effect were recorded.

## Discussion

In the present study, a preclinical study of the “Betuvax-CoV-2” vaccine was conducted to evaluate its safety and efficacy. The vaccine was tested in various toxicity studies (acute and subchronic toxicity) and, as a result, it proved safe for use. Mortality was zero, the animals gained body weight, the mass of the internal organs did not change, and no pathologies were found. Also, there was no signs of immunotoxicity or local irritation.

Reproductive toxicity test in female rats have shown that the administration of the vaccine at a dose of 5 or 20 μg or 40 μg, both before mating and during pregnancy, did not adversely affect the reproductive function or had teratogenic effects on the development of the offspring. The course of the pregnancy and pup postnatal development did not differ from the control parameters. Mortality of the pups was zero.

Multiple administration of the vaccine (7 times within a 48-day period) to male rats as part of the reproductive toxicity study did not adversely affect the pregnancy of females and the postnatal development of their offspring. Macroscopic examination of the testicles revealed no pathological changes. However, a histological study showed that male rats who received the vaccine at a dose of 20 μg or 40 μg 7 times during a 48-day period showed some change in spermatogenic epithelium. Nevertheless, Spermatogenesis Index in this group decreased insignificantly and did not affect reproductive potential of male rats. The functional state of spermatozoa did not differ from the control groups as well.

It should also be noted that subchronic toxicity study revealed no similar changes in spermatogenic epithelium, as well as other signs of reproductive toxicity in males. The changes revealed in the male reproductive toxicity study were probably associated with a high dose of the vaccine. In the study of subchronic toxicity, the drug was administered 4 times intramuscularly (1 time in 10 days) or 2 times intramuscularly (with an interval of 21 or 28 days), while in this experiment, the males received 7 intramuscular injections of the drug (once per week) at a dose of 20 μg or 40 μg, corresponding to the human dose.

The study of immunogenic properties of the “Betuvax-CoV-2” vaccine has proved its ability to stimulate both humoral and cellular immune response. Previously we showed [7] that double immunization of mice led to the formation of an adaptive T-cell immune response. We observed an increase in both CD4+ and CD8+ Tem spike-specific cells after two administrations. Moreover, based on the type of the antigen formulation (betulin adjuvant, size, and shape of nanoparticles), the Th1-directed polarization of the cytokine response with the level of the CD4+ Tem and IL-2 and TNFα was determined. A statistically significant increase was also shown for multifunctional CD8+ T cells (IFNγ+ IL-2+ TNFα+). This cytokine pattern is thought to be beneficial because patients with severe COVID-19 are usually characterized by decreased IFNγ levels and a shift to a more pronounced Th2 profile [8].

The administration of the “Betuvax-CoV-2” vaccine to mice (C57/black, BALB/c, outbred), Chinchilla rabbits, hamsters, and primates at a dose of 5 or 20 μg or 40 μg resulted in an increase in the titers of RBD-specific antibodies, specifically after the second shot. Antibodies against the RBD component of the vaccine were neutralizing, thereby providing a protective immune response against the SARS-CoV-2.

The protective activity of the vaccine was studied in golden hamsters and rhesus monkeys after being infected with the SARS-CoV-2. In a study on the suppression of the virus production in the lungs of golden hamsters, it was shown that the protection level of “Betuvax-CoV-2”, when administered intramuscularly at a dose of 20 μg or 40 μg, as well as intranasally at a dose of 4 μg or 40 μg, was 99%. Histological examination of the lungs of macaques also confirmed the findings. In the group of intramuscular injection of the vaccine, there were no pronounced signs of damage of the microvessels with developing hemorrhagic manifestations and subtotal vessel thrombosis of the pulmonary circulation as in the control group. The immunized groups did not show signs of strong intraalveolar exudation, formation of hyaline membranes, or pathological secretion in the lumen of the bronchi and bronchioles. The immunophenotypic characteristic of the infiltrate in the later stages indicated a predominant cellular immunity. In addition, immunohistochemical detection of the S-protein in the lungs of infected primates was markedly lower in the 5 µg and 20 µg intramuscular vaccine groups. It should be noted that in the study of protection activity in hamsters and macaques, the ADE effect of the vaccine was not recorded.

Thus, the results of the preclinical studies showed the immunogenicity (humoral and cell immunity), protection (significant reduction in virus loads) and safety of the vaccine, including the absence of the ADE effect, thus demonstrating a favorable “risk-benefit” ratio.

Given that there is an active development and spread of the new strains of the SARS-CoV-2, such as Omicron, a high need for frequent boosting and a use of polyvalent vaccines for different strains, that differ greatly in antigenic determinants from each other, have been preserved. The “Betuvax-CoV-2” vaccine, due to its high safety profile, efficacy, and the possibility of scalable production for different emerging strains, is well suited as a booster vaccine. We are also considering the development of an intranasal vaccine against the coronavirus, which also showed remarkable protection in this study. Currently, intramuscular injection of the “Betuvax-CoV-2” is being evaluated in a phase 1/2 of a randomized double-blind placebo-control clinical trial (NCT05270954).

## Acknowledgments

We are grateful to Murashev A.N., Rzhevsky D.I. (Branch of the Shemyakin–Ovchinnikov Institute of Bioorganic Chemistry of the Russian Academy of Sciences) for immunogenicity and safety experiments on golden hamsters and primates, Karalogly D.D. (Research Institute of Medical Primatology) for safety experiments on primates, Borisevich S.V. and Kovalchuk A. (FGBU Central Research Institute No. 48 of the Russian Ministry of Defense) for experiments of the vaccine protectiveness.

## Author Contributions

A.V.V., A.V.K., I.V.K., A.A.I. contributed to the conceptualization of experiments, A.V.V., A.V.K., N.A.K., R.V.D., A.A.N., E.O., A.V.I., E.A.R-R., M.A.S., I.V.K. — generation of data, analysis and interpretation of the results; A.V.V., N.A.K., R.V.D. — original draft preparation; A.V.V., M.E.F., K.A.B., M.D., I.V.K. — review and editing. All authors have read and agreed to the published version of the manuscript.

## Funding

The research was funded by PJSC “Human Stem Cells Institute” and Moscow Seed Fund.

## Data Availability Statement

The datasets generated or analyzed during this study are available from the corresponding author on reasonable request.

## Conflicts of Interest

A.A.I., I.V.K., M.E.F., A.V.K., A.V.V., A.V.I. have author’s rights to the patent RU2749193 “Method for preparation of betulin for use as adjuvant in vaccine against SARS-CoV-2 coronavirus” (registered by “Betuvax”). N.A.K., R.V.D., A.A.N., E.O., A.V.I., E.A.R-R., M.A.S., K.A.B., M.D. had received financial support from company “Betuvax” for provided scientific work. A.A.I., I.V.K., M.E.F., A.V.K., A.V.V. are shareholders of “Betuvax”. A.V.V. is an employee of “Betuvax”.

